# Leptin signaling and the intervertebral disc: Sex dependent effects of leptin receptor deficiency and Western diet on the spine in a type 2 diabetes mouse model

**DOI:** 10.1101/2019.12.23.887018

**Authors:** Devorah M. Natelson, Alon Lai, Divya Krishnamoorthy, Rob C. Hoy, James C. Iatridis, Svenja Illien-Jünger

## Abstract

Type 2 diabetes and obesity are associated with back pain in juveniles and adults and are implicated in intervertebral disc (IVD) degeneration. Hypercaloric Western diets are associated with both obesity and type 2 diabetes. The objective of this study was to determine if obesity and type 2 diabetes result in spinal pathology in a sex-specific manner using *in vivo* diabetic and dietary mouse models. Leptin is an appetite-regulating hormone, and its deficiency leads to polyphagia, resulting in obesity and diabetes. Leptin is also associated with IVD degeneration, and increased expression of its receptor was identified in degenerated IVDs. We used young, leptin receptor deficient (Db/Db) mice to mimic the effect of diet and diabetes on adolescents. Db/Db and Control mice were fed either Western or Control diets, and were sacrificed at 3 months of age. Db/Db mice were obese, while only female mice developed diabetes. Female Db/Db mice displayed altered IVD morphology, with increased intradiscal notochordal band area, suggesting delayed IVD cell proliferation and differentiation, rather than IVD degeneration. Motion segments from Db/Db mice exhibited increased failure risk with decreased torsional failure strength. Db/Db mice also had inferior bone quality, which was most prominent in females. We conclude that obesity and diabetes due to impaired leptin signaling contribute to pathological changes in vertebrae, as well as an immature IVD phenotype, particularly of females, suggesting a sex-dependent role of leptin in the spine.

## Introduction

Back pain is the leading cause for global disability and the most common reason for doctor visits [1]. The underlying pathologies for back pain are multifarious and are often associated with vertebral fractures, endplate defects, intervertebral disc (IVD) herniation and degeneration. Obesity is among the strongest risk factors for back pain [2,3] and also the greatest risk factor for type 2 diabetes. In the United States, obesity is rising among children and adolescents with a prevalence of 21% among those 12-19 years old [4]. The susceptibility to skeletal fractures is increased in obese children [5], and individuals with type 2 diabetes have increased fracture risk independent of bone mineral density [6]. Interestingly, bone mineral density of diabetic patients has been reported to be lower, normal, or even greater compared to age-matched non-diabetic controls [6–8], suggesting that increased fracture risk in diabetics is likely associated with alterations in bone microstructure and material properties. Body fat mass has sex dependent effects on vertebral and femoral bone quality of juveniles, demonstrated by an association of body fat mass with decreased bone stiffness [9,10]. Moreover, the prevalence of obesity and fracture risk are higher in diabetic women than men [11–13]. The literature on sex-dependent bone changes highlights a need to identify the sex effects of obesity and diabetes on vertebral pathologies in the development of spinal pathologies.

In addition to the effects on bone quality, clinical studies also demonstrated a link between obesity, type 2 diabetes, and IVD degeneration [14–17]. Yet, the evidence remains inconsistent as some studies fail to observe correlations between type 2 diabetes and IVD degeneration [18,19]. However, the hypothesis that diabetes contributes to IVD degeneration is supported by animal experiments, which demonstrate diabetes as a contributor to IVD degeneration [20]. In UCD-T2DM rats, diabetes and obesity together caused significantly decreased IVD creep strain and increased IVD stiffness, which was not observed in obese, non-diabetic control rats [21]. In mice with type 1 diabetes, we previously demonstrated decreased glycosaminoglycan content, structural deterioration in IVDs and decreased bone mass in vertebrae [22]. The literature highlights a need to identify a more mechanistic understanding of the contribution of obesity and diabetes on IVD and vertebral dysfunction in the development of spinal pathologies.

The aim of the present study was to assess the effects of type 2 diabetes and obesity on spinal health using a mouse model of diabetes (Db/Db) which develops diabetes and obesity due to leptin receptor deficiency from a point mutation within the Db gene.

Leptin is a cytokine-like hormone that primarily regulates appetite. Mice with leptin receptor deficiency exhibit polyphagia, resulting in severe obesity, elevated blood glucose, diabetes, and increased vertebral bone mass [23–25]. In addition to its role in appetite regulation leptin and its receptors have been identified in human IVDs [26,27]; and *in vitro* studies have demonstrated pro-catabolic and proinflammatory effects of leptin on nucleus pulposus (NP) and annulus fibrosus (AF) cells [28–30]. This study utilized the Db/Db genotype as well as the Western diet to enhance obesity and diabetes due to impaired leptin signaling. We hypothesized that the Db/Db mice would exhibit altered vertebral structure, IVD morphology, and spinal mechanical behavior, particularly in female mice.

## Methods

### Mouse model and experiment design

All animal experiments were performed according to the IACUC protocol at the Icahn School of Medicine at Mount Sinai, New York, NY. To investigate the effects of diet, sex and obesity-associated diabetes on spine, leptin receptor-deficient B6.BKS(D)-Lepr^db^/J (Db/Db) (15 females, 19 males) mice and their heterozygoes (Control) littermates (21 females, 27 males) mice were used (Fig 1A). After weaning, both Db/Db and Control mice were assigned to receive either Western diet (WD) or Western control diet (CD). The WD diet was formulated to mimic a “Western fast-food diet” with about 40% kcal from fat and 45% from carbohydrates (5TJN - Western Diet for Rodents, TestDiet, St. Louis, MO, USA); while the CD diet was a low-fat control for the Western diet with about 10% kcal from fat and 72% from carbohydrates (5TJS - Low Fat Control for Western Diets, TestDiet; Fig 1B). Mice were group housed with ad libitum access to water and assigned diet as well as unrestricted cage activities. At 12 weeks of age, the mice were anesthetized by Ketamine-Xylazine injection (Forane, Baxter, IL, USA) body weights were measured and mice were euthanized by cardiac puncture. After sacrifice, blood was collected for fasting blood glucose and hemoglobin A1c (HbA1c) levels. Lumbar spines were collected for structural, biochemical analysis. Coccygeal spines were used for biomechanical analyses (Fig 1A).

**Fig 1.**
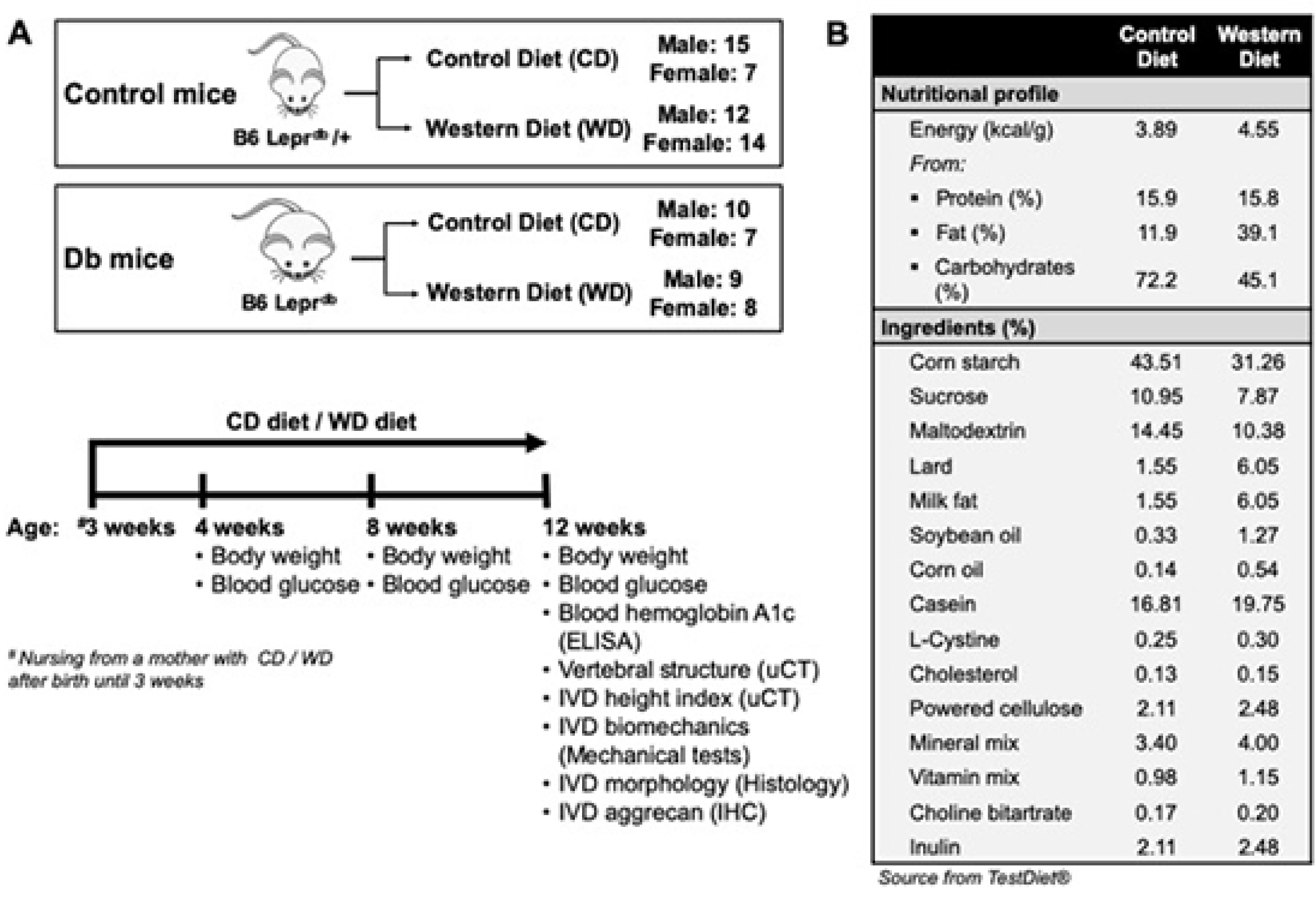
Study design. **A)** Leptin receptor deficient mice and their heterozygous controls were fed either a Western Diet or Control Diet diet. Mice were sacrificed at 12 weeks. **B)** Dietary information.

### Blood glucose and HbA1c

Fasting blood glucose and HbA1c levels were used to determine the diabetic status of the mice. After 6 hours of fasting, blood glucose was analyzed using an Aimstrip Plus Glucose Meter (Germaine Laboratories, San Antonio, TX, USA). For the measurement of HbA1c levels, the collected blood was stored in EDTA coated tubes at −80°C until further analysis. HbA1c levels were assessed using an enzymatic assay kit (Mouse Hemoglobin A1c Assay Kit #80310, Crystal Chem, Elk Grove Village, IL, USA) according to manufacturer’s instructions.

### MicroCT Assessment of Vertebral Morphology

Lumbar spines (L3-5) were dissected and stored in 10% buffered formalin phosphate fixative prior to μCT scanning. L4 vertebraes were scanned at 77-78 kV × 80 μA power with a resolution of 4.9-5.0 μm/pixel, an X-ray exposure time of 1767 ms (SkyScan 1172; Bruker Corp., Kontich, Belgium). Samples were kept hydrated in PBS while scanned in air. Hydroxyapatite phantoms (0.25 and 0.75 mg/cm3) were also scanned for mineral density calibration. μCT projections were reconstructed (N-Recon, V1.01; Bruker), and digitally aligned for consistent measurements between samples (Dataviewer V1.01; Bruker). A 1 mm region of interest was analyzed and selected to exclude growth plates based on a 0.5 mm offset from a landmark that consisted of approximately 50% of the caudal growth plate. A sole experimenter conducted all analyses to prevent bias selection of landmarks. Trabecular and cortical bone parameters were assessed following separation of bone compartments using a custom task list created in the custom processing tab of CTAn. Trabecular bone parameters assessed by 3D analysis for trabecular bone volume fraction, trabecular thickness, trabecular number, and trabecular spacing. Cortical bone parameters were assessed by 2D analysis for cortical area fraction, cross-sectional thickness, cortical area, and total area. Bone mineral density and tissue mineral density parameters were also calculated in CTAn under the Binary Image tab, keeping the threshold value consistent for all samples.

### IVD height index (DHI)

Mid-sagittal μCT images of the mice lumbar spine were used to manually identify the boundaries of the L4-5 IVD and the L4 vertebra using ImageJ. Coordinates were then used to determine IVD height and vertebral length using a customized MATLAB script (Mathworks, Inc., Natick, MA). To exclude the effects of body size between groups mice, an index was used to compare IVD heights: DHI = 2 × (DH1 + DH2 + DH3)/(A1 + A2 + A3 + B1 + B2 + B3), where A and B represent the length of the vertebral bone immediately rostral and caudal to the IVD, respectively; and DH represents the disc height between adjacent vertebrae [31].

### IVD morphology

Immediately after euthanization, L1-3 motion segments were isolated, and fixed in 10% buffered formalin. IVD-vertebrae segments were decalcified then embedded in paraffin, and sectioned sagittally at 5µm intervals for histological analysis. Mid-sagittal sections were stained with Picrosirius Red-Alcian Blue staining (PRAB) for collagen and proteoglycans, and imaged under bright field microscopy with standardized exposure time. The severity of IVD degeneration was quantified using a semi-quantitative grading system within 5 parameters for signs of degeneration, including NP structure, NP clefts/fissures, AF structure, AF clefts/fissures, and NP/AF boundary [32]. All sections were examined by two researchers blinded to the experimental groups, and then averaged for analysis [33]. The boundaries of notochordal band and NP were manually defined in photoshop, and the area of notochordal band relative to area of NP was calculated by outlining both notochordal band and total NP area in photoshop. IVD cellularity was visualized using hematoxylin staining with eosin as counter-staining [34], and then imaged under bright field microscopy with standardized exposure time. IVD cells within the notochordal band were manually counted using ImageJ, and then normalized to the area counted.

### Motion segment biomechanics

Biomechanical properties of caudal vertebra-IVD-vertebra motion segment (Co4-5) were assessed via tension-compression, creep, and torsional tests; which provide substantial information to evaluate properties of NP pressurization and hydration, AF lamellae integrity and quality, as well as IVD laxity and viscoelasticity. Immediately after dissection, Co4-5 motion segments were wrapped in phosphate buffer saline (PBS)-soaked paper towels and stored at −80°C until the day of testing. Axial and creep tests were performed using the ElectroForce 3200 testing machine (Bose Corporation, Eden Prairie, MN), and torsional tests were performed using the AR2000x Rheometer (TA Instruments, New Castle, DE, USA).

The testing protocol was adopted from our previous studies [35–37]. In brief, on the day of testing, after 10 mins in PBS for thawing and hydrating, the motion segment specimens were loaded into parallel-platens of an axial loading machine (Bose ElectroForce 3220; TA Instruments, New Castle, DE, USA) using a custom-designed fixture with a fluid bath of PBS for axial testing. During the axial testing, specimens underwent 20 cycles of ±0.5 N tension-compression test at 1 Hz, followed by 1 min of dwelling to allow the specimens to relax, and then 45 mins of creep test with the compressive force at 0.5 N. After 30 min of rehydration in PBS, specimens were attached to a rheometer (AR2000; TA Instruments, New Castle, DE, USA) using a custom-designed fixture for torsional testing which consisted of 20 cycles of ±10° rotation in both directions at 1 Hz, followed by torsion-to-failure test at the rate of 1°/s. The loading profile obtained from the 20th cycle of tension-compression and torsional tests were used to determine compressive stiffness, tensile stiffness, axial range of motion, axial neutral zone length, torsional stiffness (average from the stiffness obtained from clockwise and counterclockwise direction), torsional neutral zone length and torque range using custom-written MATLAB codes. For analyzing the characteristics of creep test, a 5-parameter viscoelastic solid model was applied to calculate creep and total displacements, elastic stiffness, time constant (*τ*) and stiffness for both fast response and slow response using a custom-written MATLAB code. For the torsion-to-failure test, the failure strength and angle to failure were identified manually from the loading profile.

### Statistical analyses

Body weight, blood glucose level, HbA1c, vertebral morphology, DHI, IVD biomechanics, and IVD score were compared using two-way ANOVA to assess the effects of genotype and diet. Tukey’s post-hoc comparison was performed to assess the effects of Western diet and Db/Db genotype on vertebral and IVD changes. Results were analyzed for females and males separately and displayed as average ± standard deviation. All statistical analyses were performed using Graphpad Prism7 (GraphPad Software, Inc., La Jolla, CA) with level of significance set at 0.05.

## Results

### General observations confirmed an obese and pre-diabetic phenotype

As expected, at 3 months, Db/Db genotype and WD caused significantly increased body weight (Fig 2). Db/Db genotype caused significantly increased HbA1c levels and was highest in female Db/Db WD mice (Fig 3). Blood glucose was significantly increased in female mice with Db/Db genotype at all time points and levels were greatest for Db/Db WD mice (Fig 3). In males, blood glucose was significantly increased with WD at 4 & 8 weeks but not at 12 weeks. Overall, HbA1c results indicate that all Db/Db mice were pre-diabetic and female Db/Db WD mice were diabetic [38].

**Fig 2.**
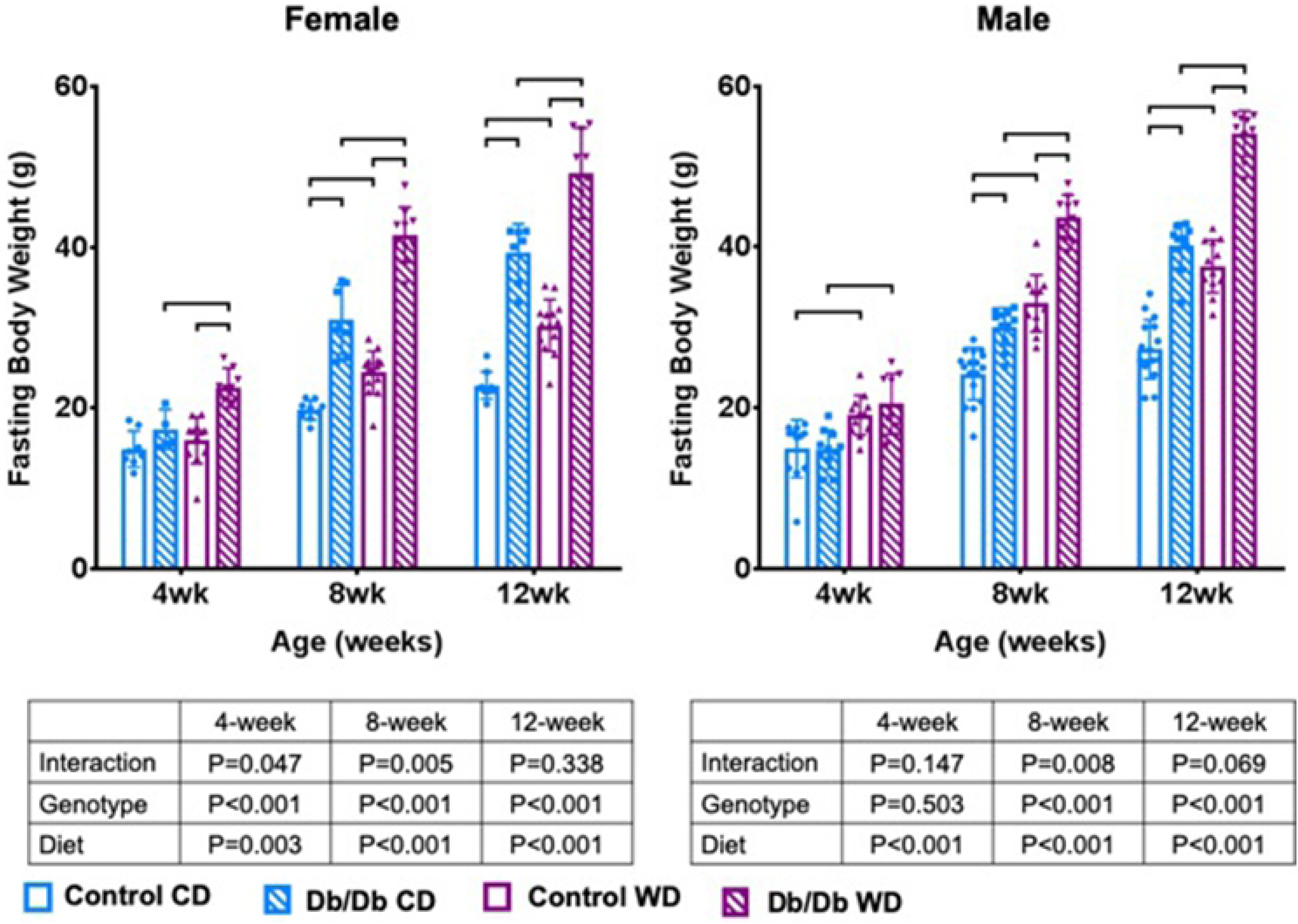
Leptin receptor deficiency and Western Diet both increased fasting body weight in mice, for both male and female groups. p < 0.5. wk=weeks.

**Fig 3.**
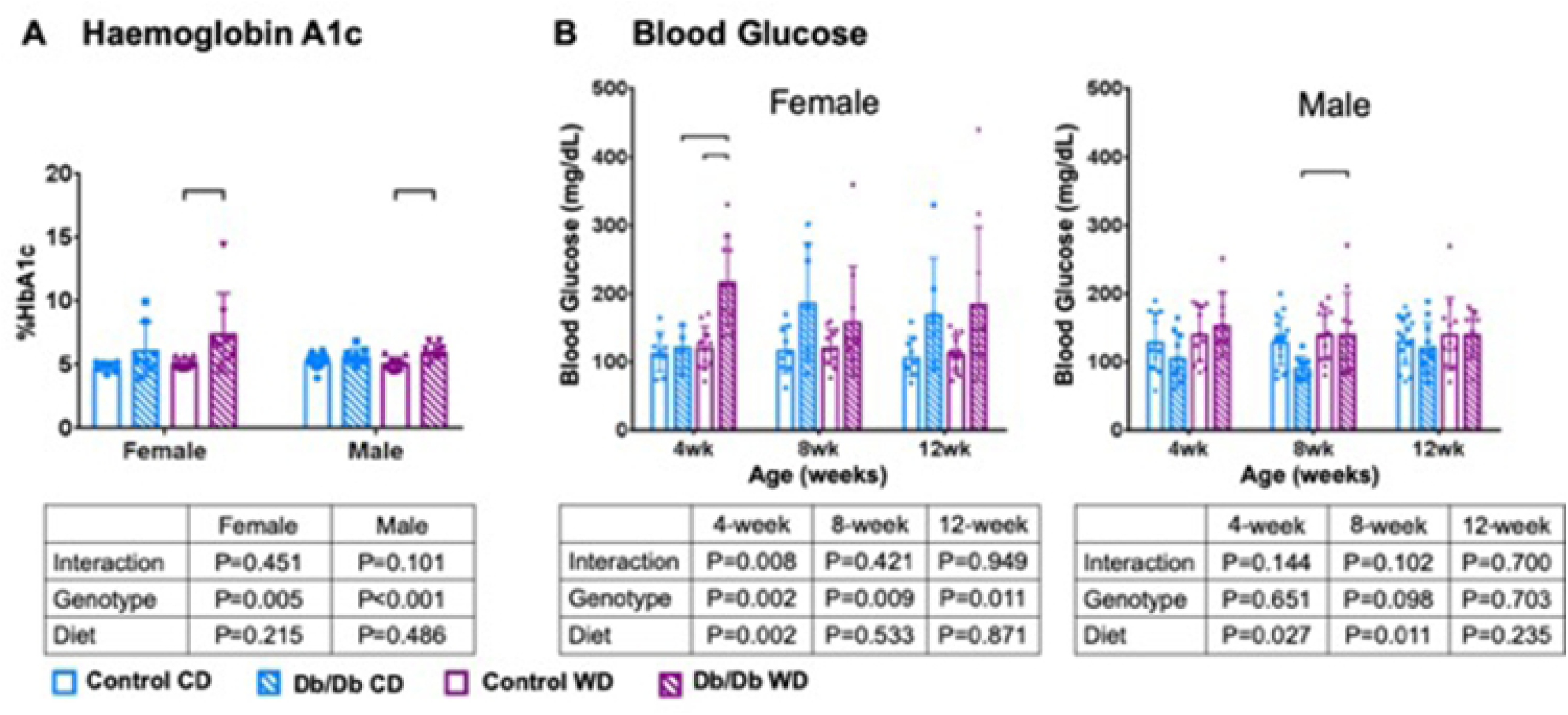
Leptin receptor deficiency increased HbA1c in mice for both male and female groups. **A)** HbA1c was elevated to diabetic levels in female mice. **B)** Blood glucose did not reach diabetic levels. p < 0.5. wk=weeks.

### Western diet and Db/Db genotype had sex-dependent effects on vertebral bone microstructure

Trabecular bone structure was strongly affected by Db/Db genotype. Both female and male Db/Db mice had increased trabecular bone volume fraction with increased trabecular number and decreased trabecular spacing, which was independent from diet (Fig 4, Table 1). In contrast, only female Db/Db mice on WD had decreased cortical bone volume fraction, cortical thickness, and cortical area compared to female WT mice on WD. No differences were observed in male mice for any cortical parameter; suggesting that changes in bone structure due to Db/Db genotype were sex-dependant (Fig 5, Table 1).

**Table 1.** uCT parameters.

|  |  | Female |  |  |  |  |  | Male |  |  |  |  |  | LF versus HF (*p-value*) |  |  |  |
| --- | --- | --- | --- | --- | --- | --- | --- | --- | --- | --- | --- | --- | --- | --- | --- | --- | --- |
|  |  | LF |  | *p* | HF |  | *p* | LF |  | *p* | HF |  | *p* | Female |  | Male |  |
|  |  | Control | Db/Db |  | Control | Db/Db |  | Control | Db/Db |  | Control | Db/Db |  | Control | Db/Db | Control | Db/Db |
| Trabecular | Tb. BS/BV (mm<sup>-1</sup>) | 69.53<br>±2.59 | 66.02<br>±7.41 | 0.625 | 68.11<br>±4.38 | 75.48<br>±6.26 | **0.03** | 75.8<br>±5.82 | 70.25<br>±5.51 | 0.458 | 76.93<br>±6.08 | 75.54<br>±11.44 | 0.981 | 0.951 | **0.009** | 0.989 | 0.498 |
| Cortical | Tt.Ar (mm<sup>2</sup>) | 1.47<br>±0.08 | 1.56<br>±0.07 | 0.999 | 1.46<br>±0.14 | 1.45<br>±0.09 | 0.115 | 1.61<br>±0.09 | 1.68<br>±0.13 | 0.337 | 1.516<br>±0.12 | 1.654<br>±0.10 | 0.961 | 0.278 | 0.997 | 0.492 | 0.062 |
|  | Ct. BS/BV (mm<sup>-1</sup>) | 41.50<br>±3.23 | 46.17<br>±3.97 | 0.167 | 38.53<br>±2.74 | 50.14<br>±5.73 | **<0.001** | 42.65<br>±3.17 | 46.87<br>±1.83 | **0.037** | 41.32<br>±2.32 | 45.25<br>±4.21 | **0.049** | 0.468 | 0.243 | 0.794 | 0.697 |

**Fig 4.**
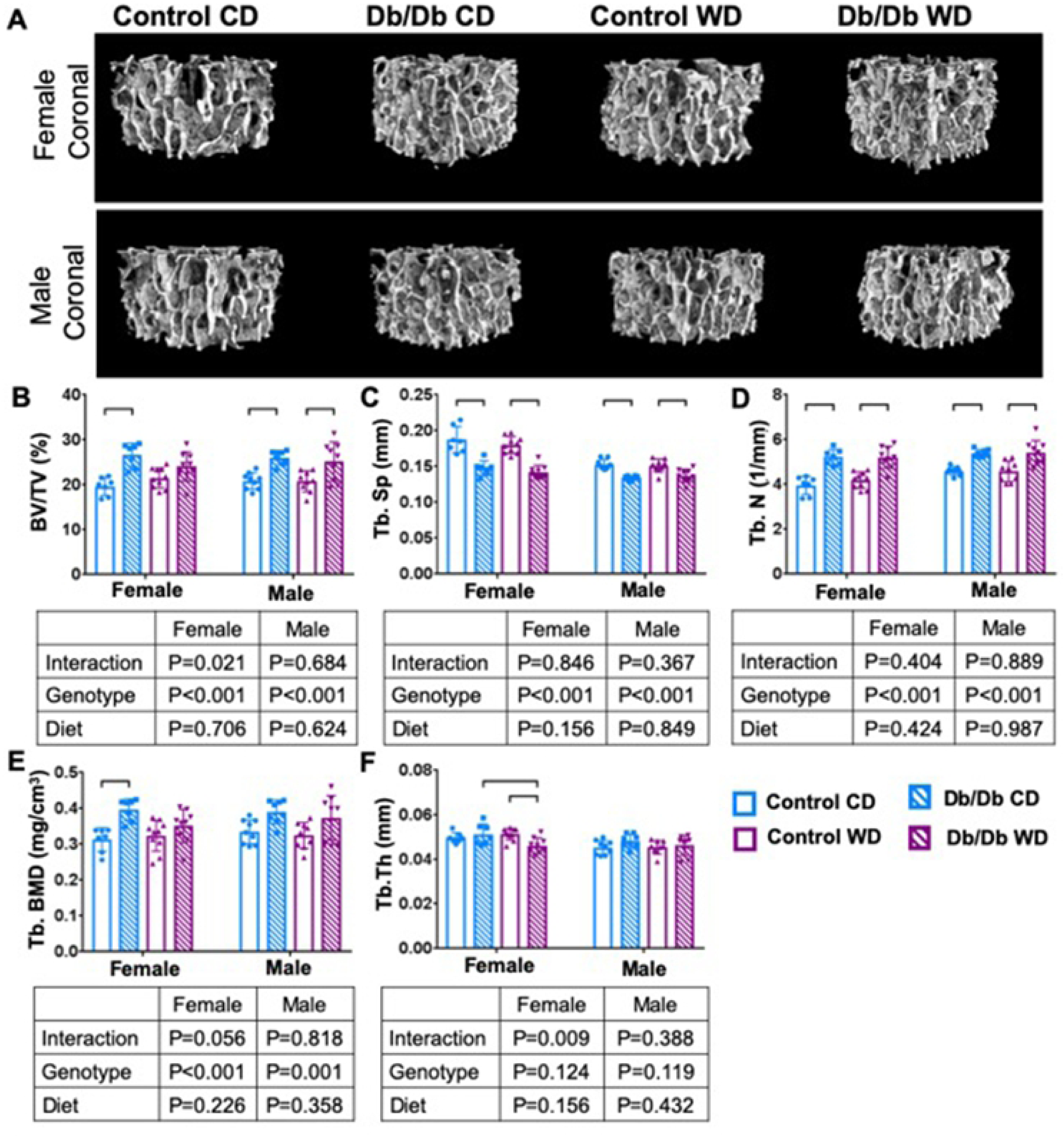
Trabecular bone increased with leptin receptor deficiency. **A)** 3D µCT images of trabecular bone of (top) female and (bottom) male mice. **B)** BV/TV C) Tb.Sp, and **D)** Tb.N demonstrate inferior Tb microstructure with Db/Db genotype. Only female Db/Db mice had **E)** increased Tb. BMD and **F)** decreased Tb.Th. p < 0.05. BV/TV = bone volume fraction; Tb.Sp = trabecular spacing; Tb.N – trabecular number; Tb. BMD = trabecular bone mineral density; Tb.Th = trabecular thickness.

**Fig 5.**
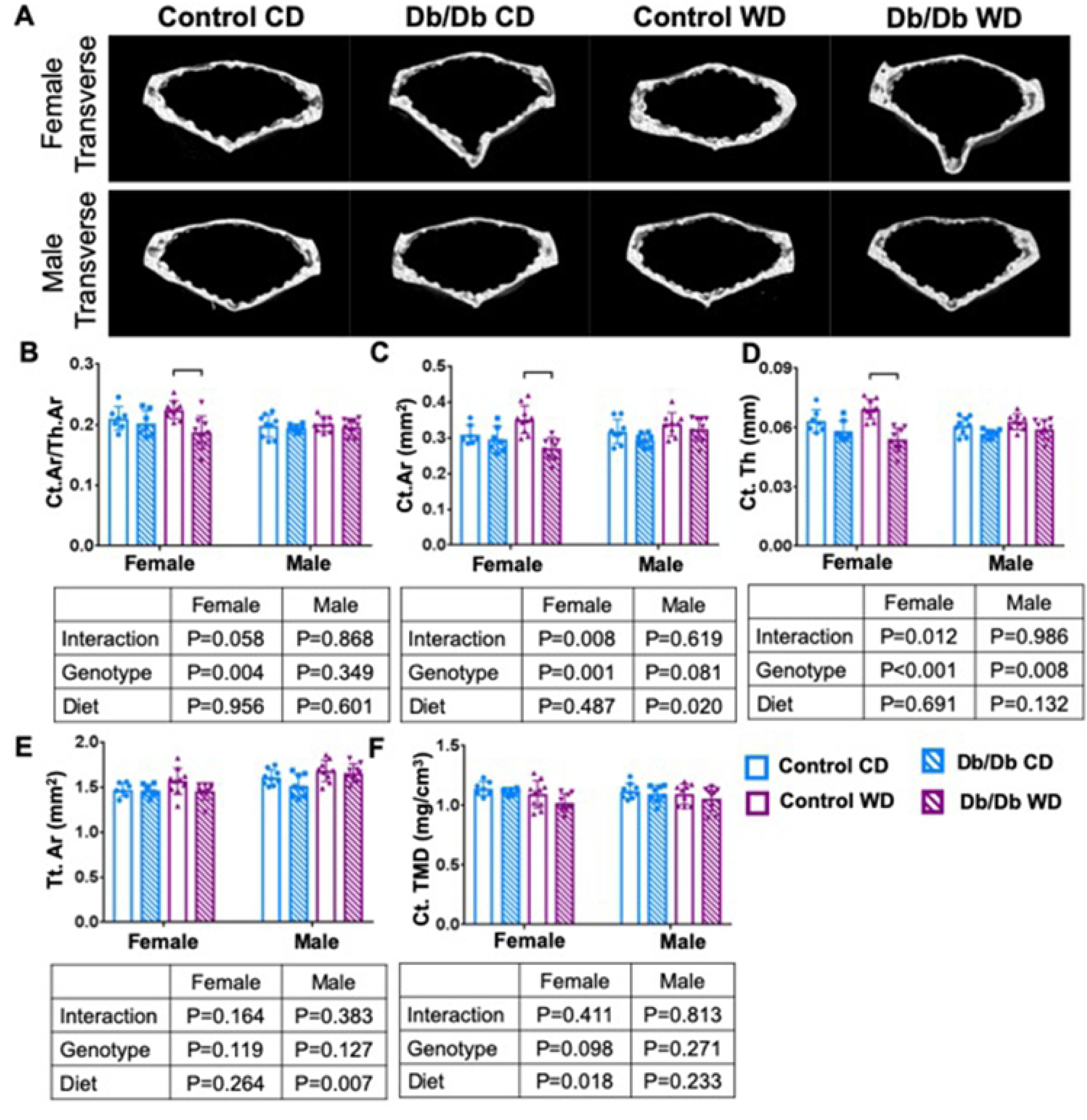
Leptin receptor deficiency caused decreased in cortical bone for female mice only. A) 3D µCT images of cortical bone for (top) female and (bottom) male mice. Leptin receptor deficiency decreased B) Ct.Ar/Th.Ar C) Ct.Ar and D) Ct.Th. No changes were observed for E) Tt.Ar or F) Ct.TMD. Male mice showed not changes any parameter. p < 0.05. Ct.Ar/Th.Ar = cortical area fraction; Ct.Ar = cortical area; Ct.Th = cortical thickness; Tt.Ar = total area; Ct. TMD cortical tissue mineral density.

### Db/Db genotype caused sex dependent changes in IVD morphology, but did not induce IVD degeneration

Pronounced differences in IVD morphology were observed between female Control and Db/Db mice (Fig 6). IVDs of female Db/Db mice had a significantly increased notochordal band area with large, vacuolated cells, and significantly fewer cells per area compared to female control mice. These changes were independent from diet. Despite these pronounced differences in morphology, leptin receptor deficiency did not cause IVD degeneration in 3 months old mice. There was no difference in Tam IVD grading score between any groups. No effects of diet or genotype on IVD structure or morphology were detected in male mice. No differences in DHI was observed for any group, although some significant but subtle alterations in vertebral length and IVD height were identified (Fig 7).

**Fig 6.**
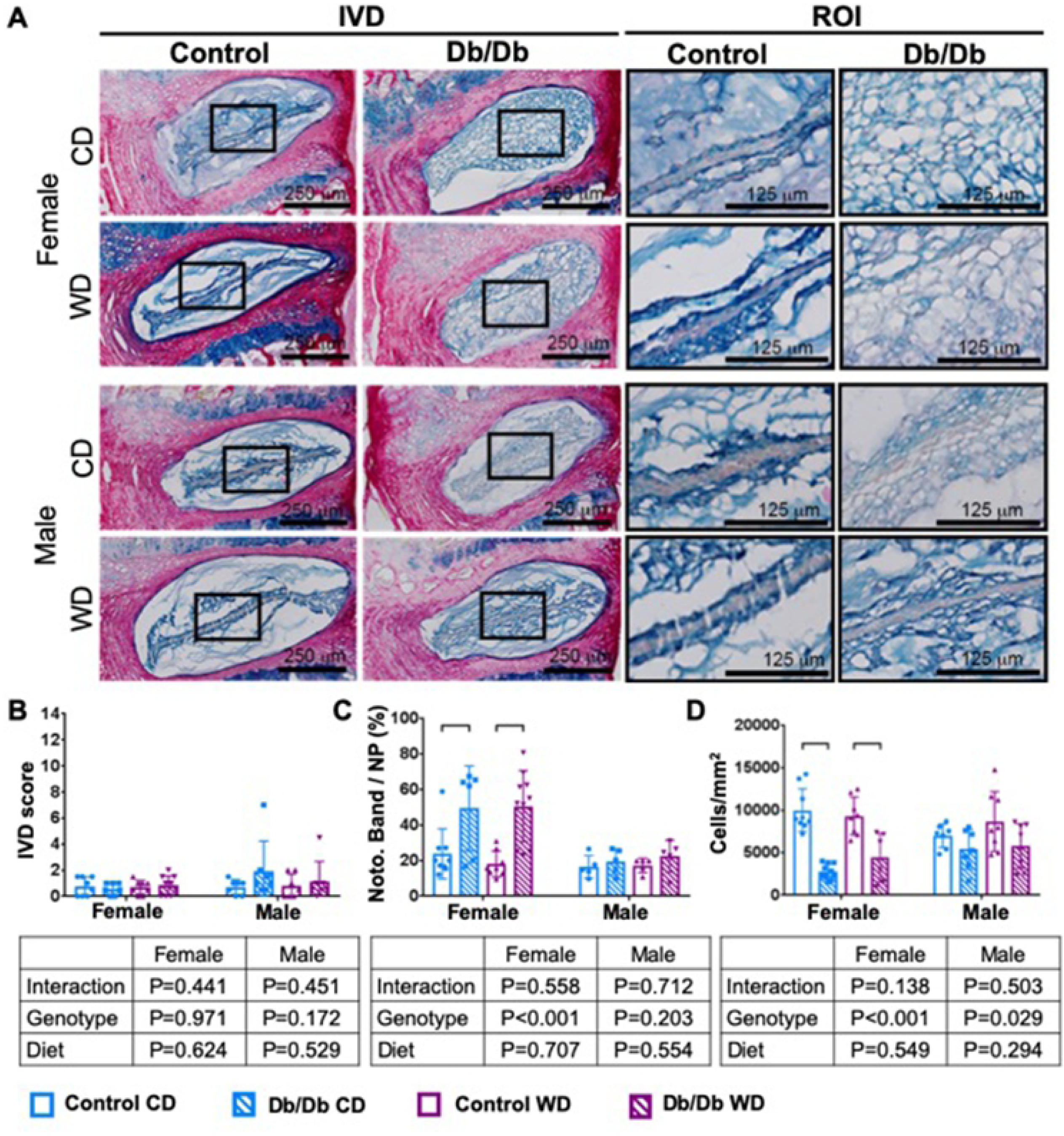
Notochordal band size increased in female leptin receptor deficient mice. **A)** Representative Picrosirius red/Alcian blue images demonstrate an increased notochordal band in (top) female Db/Db mice. **B)** Db/Db genotype did not cause IVD degeneration. In female Db mice, **C)** notochordal cell size was increased and **D)** cells per area was decreased. No changes were observed in male IVDs. Black boxes mark region of interest (ROI).

**Fig 7.**
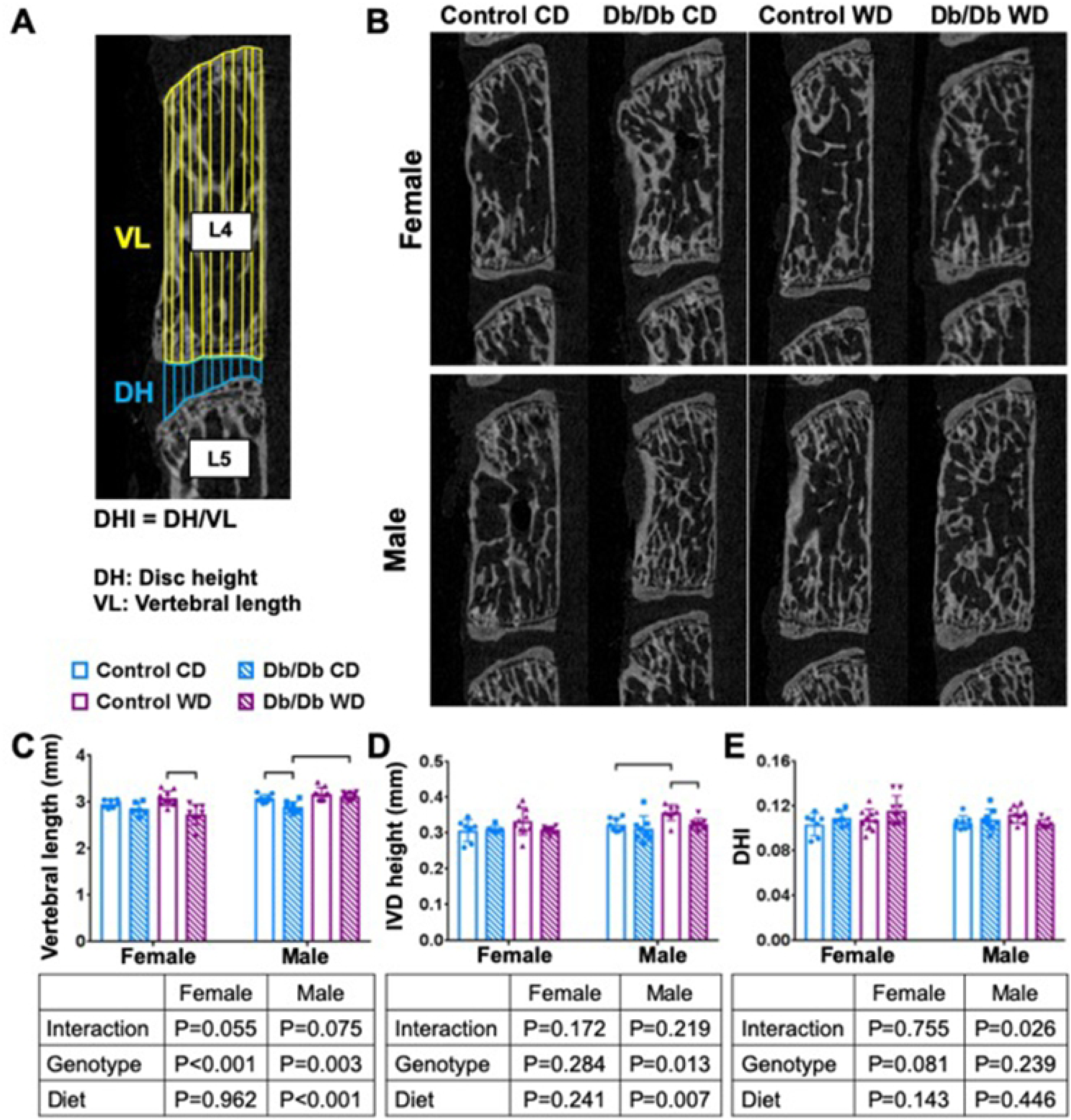
Leptin receptor deficiency caused decrease vertebral length and disc height, but no change in disc height index. **A)** Schematic of IVD height and vertebrae length measurement. **B)** µCT midsagittal sections of vertebral bone for (top) female and (bottom) male. p < 0.05. **C)** vertebral length **D)** IVD height **E)** DHI.

### IVD torsional, but not axial, biomechanical behavior was compromised with leptin receptor deficiency

Leptin receptor deficiency significantly diminished torsional biomechanical properties of mice caudal motion segments. In Db/Db mice, cyclic torsional testing showed significantly decreased torsional stiffness and the torsion-to-failure test further demonstrated significantly decreased torsional failure strength with Db/Db genotype for both sexes. WD caused a small but significant increase in torsional failure strength in both sexes. The angle-to-failure, on the other hand, significantly decreased in female Db/Db mice only, while no changes were shown in male Db/Db mice. It should be noted that the majority of motion segments failed at the growth plate, suggesting a weakness in Db/Db vertebral bones which were more greatly affected for females (Fig 8, Table 2).

**Table 2.** Biomechanical properties.

|  |  | Female |  |  |  |  |  | Male |  |  |  |  |  | LF versus HF (*p*) |  |  |  |
| --- | --- | --- | --- | --- | --- | --- | --- | --- | --- | --- | --- | --- | --- | --- | --- | --- | --- |
|  |  | LF |  |  | HF |  |  | LF |  |  | HF |  |  | Female |  | Male |  |
|  |  | Control | Db/Db | *p* | Control | Db/Db | *p* | Control | Db/Db | *p* | Control | Db/Db | *p* | Control | Db/Db | Control | Db/Db |
| Axial | Compressive Stiffness (N/mm) | 16.59<br>±4.98 | 14.32<br>±3.65 | 0.797 | 15.68<br>±3.09 | 17.38<br>±3.68 | 0.844 | 19.05<br>±5.08 | 18.95<br>±3.81 | >0.99 | 18.83<br>±7.42 | 17.32<br>±4.54 | 0.934 | 0.970 | 0.613 | >0.999 | 0.906 |
|  | Tensile Stiffness (N/mm) | 9.15<br>±3.89 | 12.10<br>±4.10 | 0.705 | 11.61 ±4.12 | 13.31<br>±6.77 | 0.858 | 12.81<br>±4.17 | 12.06<br>±4.40 | 0.976 | 11.92<br>±4.15 | 11.84<br>±3.52 | >0.99 | 0.719 | 0.965 | 0.967 | 0.999 |
|  | Range of Motion (mm) | 0.19<br>±0.04 | 0.19<br>±0.03 | >0.99 | 0.17<br>±0.02 | 0.20<br>±0.03 | 0.216 | 0.17<br>±0.04 | 0.18<br>±0.05 | 0.895 | 0.16<br>±0.04 | 0.18<br>±0.04 | 0.694 | 0.667 | 0.846 | 0.985 | >0.999 |
|  | Neutral Zone Length (mm) | 0.07<br>±0.03 | 0.05<br>±0.02 | 0.419 | 0.06<br>±0.02 | 0.07<br>±0.02 | 0.525 | 0.06<br>±0.02 | 0.07<br>±0.03 | 0.543 | 0.06<br>±0.02 | 0.07<br>±0.02 | 0.974 | 0.787 | 0.254 | 0.956 | 0.984 |
|  | τ1 (slow response) (sec) | 51.71<br>±27.13 | 54.75<br>±12.16 | 0.992 | 41.85<br>±13.84 | 37.68<br>±16.55 | 0.965 | 45.13<br>±9.83 | 41.79<br>±21.00 | 0.975 | 32.51<br>±14.78 | 53.22<br>±20.41 | 0.086 | 0.683 | 0.392 | 0.449 | 0.487 |
|  | τ2 (slow response) (sec) | 739.0<br>±255.8 | 900.8<br>±92.1 | 0.57 | 707.7<br>±242.7 | 669.0<br>±194.4 | 0.982 | 724.7<br>±182.8 | 746.2<br>±134.8 | 0.989 | 453.1<br>±37.8 | 806.0<br>±171.8 | **0.001** | 0.990 | 0.268 | **0.009** | 0.821 |
|  | Creep Displacement (mm) | 0.20<br>±0.05 | 0.21<br>±0.02 | 0.972 | 0.18<br>±0.03 | 0.18<br>±0.07 | 0.999 | 0.19<br>±0.05 | 0.17<br>±0.05 | 0.679 | 0.15<br>±0.04 | 0.19<br>±0.04 | 0.244 | 0.853 | 0.624 | 0.170 | 0.801 |
| Creep | Total Displacement (mm) | 0.24<br>±0.06 | 0.25<br>±0.03 | 0.982 | 0.22<br>±0.04 | 0.22<br>±0.07 | 0.994 | 0.23<br>±0.06 | 0.20<br>±0.05 | 0.598 | 0.18<br>±0.03 | 0.23<br>±0.04 | 0.257 | 0.882 | 0.635 | 0.186 | 0.718 |
|  | Elastic Stiffness (N/mm) | 11.66<br>±2.47 | 12.56<br>±2.39 | 0.950 | 13.06<br>±3.11 | 13.08<br>±3.18 | >0.999 | 14.65<br>±4.19 | 16.57<br>±3.91 | 0.713 | 16.03<br>±4.53 | 14.12<br>±3.02 | 0.752 | 0.757 | 0.990 | 0.888 | 0.534 |
|  | Stiffness1 (N/mm) | 12.64<br>±2.00 | 14.94<br>±2.73 | 0.706 | 17.53<br>±4.44 | 15.10<br>±3.43 | 0.548 | 17.01<br>±6.96 | 19.44<br>±9.51 | 0.917 | 27.17<br>±10.17 | 17.22<br>±5.31 | 0.079 | 0.059 | >0.999 | 0.071 | 0.935 |
|  | Stiffness2 (N/mm) | 3.154<br>±0.94 | 2.75<br>±0.31 | 0.747 | 3.31<br>±0.50 | 3.12<br>±0.79 | 0.943 | 3.27<br>±0.93 | 3.73<br>±0.93 | 0.664 | 4.08<br>±0.97 | 2.99<br>±0.89 | 0.078 | 0.965 | 0.810 | 0.243 | 0.294 |
| Torsion | Torque Range (Nm) | 0.003<br>±0.001 | 0.003<br>±0.001 | 0.980 | 0.004<br>±0.002 | 0.003<br>±0.002 | 0.432 | 0.003<br>±0.002 | 0.003<br>±0.001 | 0.667 | 0.004<br>±0.002 | 0.003<br>±0.001 | 0.343 | 0.978 | 0.945 | 0.975 | 0.997 |
|  | Neutral Zone Length (rad) | 0.39<br>±0.04 | 0.37<br>±0.03 | 0.916 | 0.39<br>±0.04 | 0.40 ± 0.07 | 0.946 | 0.34 ± 0.06 | 0.38 ± 0.08 | 0.414 | 0.36 ± 0.07 | 0.39<br>±0.04 | 0.675 | 1.000 | 0.652 | 0.962 | 1.000 |

**Fig 8.**
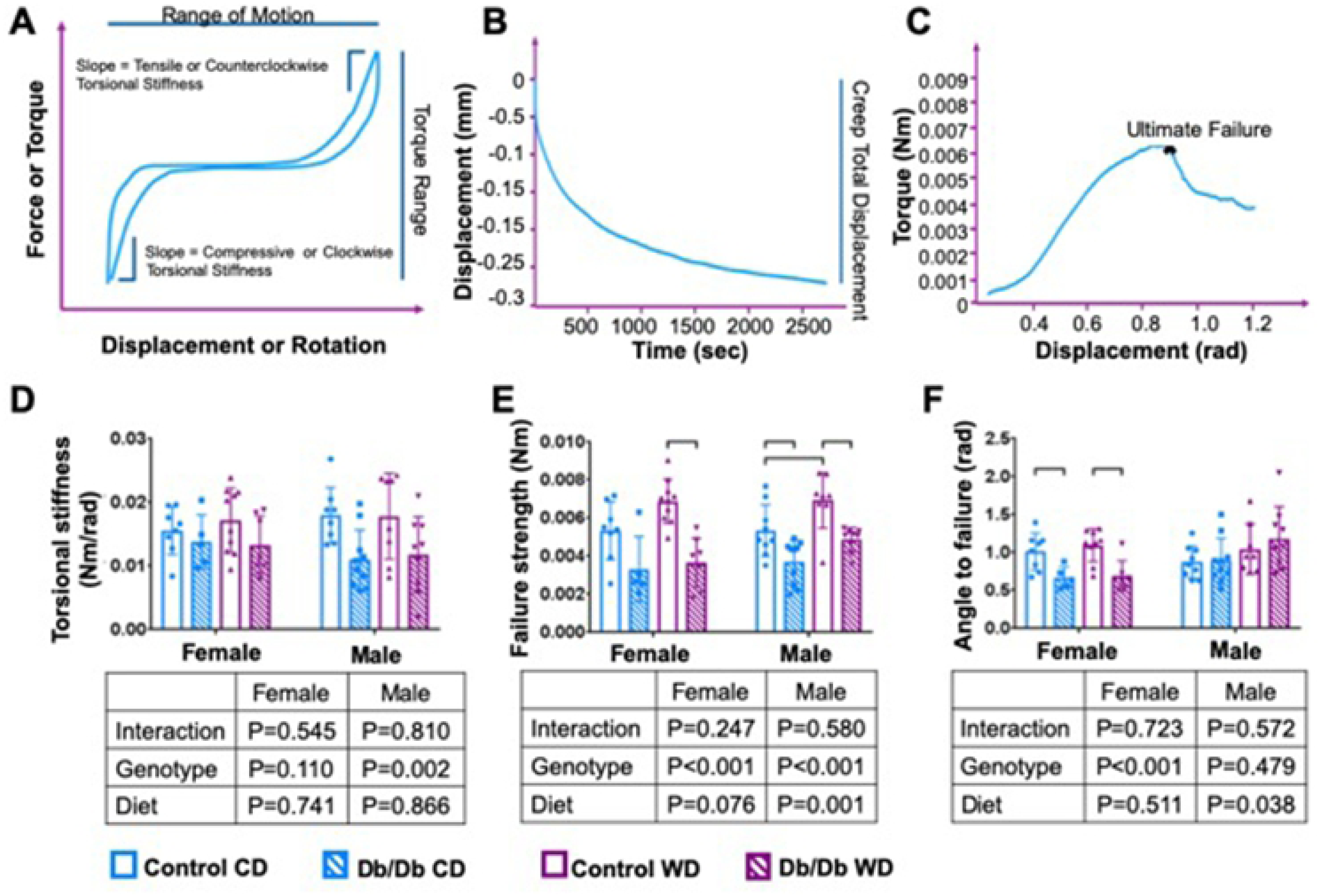
Torsional strength decreased with Db/Db genotype. Schematic curves from **A)** axial-compression and torsion testing, **B)** Creep, and **C) t**orsion to failure curves after biomechanical testing analysis. **D)** torsional stiffness **E)** Failure strength **F)** angle to failure. p<0.05

There were no significant differences between groups in axial biomechanical properties. Neither compressive and tensile stiffness, nor range of motion, or axial neutral zone length showed any significant changes associated with Db/Db genotype or WD. Additionally, there were no changes in creep parameters with Db/Db genotype or WD. No differences were observed in fast or slow time constants, creep displacements, or stiffnesses. Together, the highly significant changes in torsional biomechanical properties suggests leptin receptor deficiency and obesity had the largest functional changes in vertebral properties.

## Discussion

There is a need for a more mechanistic understanding of interactions between obesity, type 2 diabetes and spinal pathologies because of their increasing prevalence. This study used Db/Db mice with leptin receptor deficiency and Western diet to test the hypothesis that type 2 diabetes and obesity would develop dysfunctional vertebral structure, IVD morphology, and spinal biomechanical function in a sex-dependent manner. Db/Db mice and Western Diet both resulted in severe obesity, and Db/Db genotype also significantly increased HbA1c indicating diabetic/prediabetic conditions. Db/Db genotype resulted in the severe alterations in spinal structures that were most prominent in females and subtle or not present in males. Specifically, Db/Db genotype increased vertebral trabecular bone volume fraction and density, decreased vertebral cortical thickness and area, disrupted IVD morphology, and resulted in increased motion segment torsional failure risk. Interestingly, Db/Db mice did not develop any signs of IVD degeneration and IVDs instead appeared less mature with large notochordal cells populating the NP, suggesting that diabetes and impaired leptin signaling had more substantial effects on vertebrae than on IVDs. While some significant effects of WD were detected on spinal structures, the Db/Db genotype had the most severe effects, indicating that pre-diabetic/diabetic conditions had larger effects than obesity conditions on spinal structures.

The most important finding of this study is that Db/Db genotype caused increased vertebral trabecular bone density, decreased cortical bone quality, and decreased motion segment torsional failure strength. Clinically, obesity and type 2 diabetes are often associated with increased bone mineral density and fracture risk [39–41], which has been speculated to involve reduced cortical bone density in type 2 diabetics [42,43]. In this study, Db/Db genotype resulted in the most substantial changes in bone with the two-way ANOVA detecting significantly increased trabecular BV/TV and BMD in both males and females that involved increased trabecular number and decreased trabecular spacing. No significant effects of diet were detected on trabecular bone properties, suggesting this was predominantly a diabetic phenotype and not an obesity phenotype. The Db/Db genotype also significantly diminished cortical structure for both sexes in the two-way ANOVA, although female Db/Db mice on WD had the most significant cortical changes. Db/Db genotype also significantly reduced torsional failure strength for both sexes indicating that these bone structural changes caused significant biomechanical dysfunction. In contrast, WD significantly increased failure strength on two-way ANOVA, leading us to conclude that the increased fracture risk involved the pre-diabetic/diabetic condition and not obesity. Similar to our study, Williams et al. found that, lumbar vertebrae of 10 week-old male Db/Db mice had decreased cortical thickness but no changes in trabecular bone volume fraction [44]. Huang et al. used lumbar vertebrae of 36 week old male mice and demonstrated that leptin receptor deficiency increased bone volume fraction and density and increased trabecular number [45], also consistent with our findings. Together these studies amplify the need to investigate cortical and trabecular bone compartments distinctly when assessing vertebral bone quality, with implications that cortical bone structure most likely plays a critical role in spinal torsional failure risk. We therefore conclude that pre-diabetic/diabetic conditions from leptin receptor deficiency play a significant role in dysfunctional vertebral structure that can increase fracture risk, and these findings have important parallels with the clinical observations.

Female Db/Db mice on WD had the most substantial effects on vertebral bone structure and IVDs. The association of metabolic syndrome with low back pain has greater prevalence in women than in men [46]. Sex dependent effects of leptin receptor deficiency were also suggested in a recent study by McCabe et al. [47] who demonstrated that alterations of specific leptin receptor sites contributed to sex-dependent bone responses to leptin, which could be particularly relevant during juvenile obesity, where loss of leptin signaling could diminish bone development and growth [47]. Sex hormones such as sex hormone-binding globulin have been identified to have sex-dependent effects on leptin and Type 2 diabetes [48,49], and might explain the sex-dependent effects of leptin that were observed in this study. Future studies are required to address the effects of sex-hormones on leptin receptor deficiency mechanistically. We conclude that female mice had the most severe effects of Db/Db genotype and WD on spinal structures; although we note that female on WD had the most significant elevation of HbA1c levels, suggesting that their larger alterations in spinal structures may also be due to the greatest severity of the diabetic condition.

The lack of degenerative changes in IVDs of Db/Db mice is in contrast with our previous studies on western style diets in C57BL/6J mice, which demonstrated that pre-diabetes contributed to inferior vertebral quality, accelerated progression of IVD degeneration, and increased leptin levels [50]. Leptin is increased in degenerated IVDs [26] and thought to stimulate cell proliferation of human NP cells via its receptors OBRa and OBRb [51]. Taking into account its contribution to IVD degeneration in previous studies, diminished leptin signaling of this study might have been protective against IVD degeneration by maintaining large vacuolated cells in the notochordal band via decelerating NP cell proliferation and differentiation. Based on these observations, we speculate that leptin deficiency may have decelerated IVD degeneration despite the increased body weight and pre-diabetic status which caused IVD degeneration in prior animal models. Nevertheless, longer duration studies and aging may accumulate more IVD structural alterations suggestive of degeneration, as was observed in our type 1 diabetes model [22].

This study applied Db/Db mice and WD in order to distinguish obesity effects from type 2 diabetes effects in adolescent mice. Highly processed Western diets cause excess calorie intake and are associated with the rising prevalence of obesity [52] and diabetes [52]. Our diet resembled a Western diet that is high in fat and low in carbohydrates and caused increased weight gain all groups; however, effects on spinal structures were overshadowed by the Db/Db genotype. While Db/Db genotype significantly increased HbA1c and blood glucose levels at some point during the study period, not all our Db/Db mice developed sustained hyperglycemia. Hyperglycemia in B6.BKS(D)-Lepr^db^/J is transient and does not manifest with stable elevated levels in all Db/Db mice [53]. To fully assess the diabetic status, we assessed HbA1c levels in addition to hyperglycemia, which revealed that most female Db/Db mice on WD did develop diabetes at some time during their lifespan and many mice in the Db/Db cohorts exhibited pre-diabetic conditions. The Db/Db model is well accepted in the literature as a type 2 diabetes model, although other diabetic models exist and assessments of spinal pathologies in additional diabetic mouse models and/or leptin impairments [54] would enable broadened interpretations. This model demonstrated that prediabetes/diabetes resulted in more substantial changes in spinal structures than obesity in these adolescent mice, although some effects of obesity were detected and longer duration diets in older mice are likely to result in more substantial spinal changes in support of the clinical literature that obesity can induce spinal changes in juveniles and adults [15,55].

In conclusion, prediabetes/diabetes from leptin receptor deficiency resulted in severe cortical and trabecular bone changes and diminished torsional failure strength. Pre-diabetes/diabetes dominated obesity effects and these changes were greatest in females. No evidence for IVD degeneration was observed, and taken with the literature, results suggested a potential protective role of impaired leptin signaling against diabetes- and obesity-induced IVD degeneration.

## Significance

The tremendous public health burden of back pain is increasing, and is likely to grow due to known associations with obesity and DM, which are also increasing in prevalence. This study directly informs physiological factors important in spinal health, emphasizes the need to investigate sex differences, and highlights a potential role for leptin in the development and maturation of the spine.

## Acknowledgements

The authors thank Damien M. Laudier and Madeline P. Smith for their technical assistance.

